# Multiethnic Polygenic Risk Prediction in Diverse Populations through Transfer Learning

**DOI:** 10.1101/2022.03.30.486333

**Authors:** Peixin Tian, Tsai Hor Chan, Yong-Fei Wang, Wanling Yang, Guosheng Yin, Yan Dora Zhang

## Abstract

Polygenic risk scores (PRS) leverage the genetic contribution of an individual’s genotype to a complex trait by estimating disease risk. Traditional PRS prediction methods are predominantly for European population. The accuracy of PRS prediction in non-European populations is diminished due to much smaller sample size of genome-wide association studies (GWAS). In this article, we introduced a novel method to construct PRS for non-European populations, abbreviated as TL-Multi, by conducting transfer learning framework to learn useful knowledge from European population to correct the bias for non-European populations. We considered non-European GWAS data as the target data and European GWAS data as the informative auxiliary data. TL-Multi borrows useful information from the auxiliary data to improve the learning accuracy of the target data while preserving the efficiency and accuracy. To demonstrate the practical applicability of the proposed method, we applied TL-Multi to predict the risk of systemic lupus erythematosus (SLE) in Asian population and the risk of asthma in Indian population by borrowing information from European population. TL-Multi achieved better prediction accuracy than the competing methods including Lassosum and meta-analysis in both simulations and real applications.

## 1 INTRODUCTION

Genetic risk prediction is an important methodology for understanding the underlying genetic architecture and the inclusion of information on complex traits, such as estimating the genetic risk of complex traits or diseases (e.g., coronary artery disease)(Chatterjee et al., 2016; Ge et al., 2019). Polygenic risk scores (PRS) are one of the approaches to reflect a mathematical aggregation of risk by variants such as single nucleotide polymorphisms (SNPs) (Peterson et al., 2019). With the application of the best linear unbiased predictor to estimate PRS, some methods use summary association statistics as training data (Consortium, 2009; Vilhjálmsson et al., 2015; Shi et al., 2016), and others require individual-level data, such as genotype data and phenotypes (De Los Campos et al., 2010; Speed and Balding, 2014; Maier et al., 2015; Moser et al., 2015; Coram et al., 2017). As an implementation, PRS have become a widely used statistical tool to estimate the genetic risk of certain diseases or phenotypes (Mak et al., 2017). Specifically, PRS for a particular disease demonstrate the risk index for people to suffer from the disease. A remarkable study of five common diseases (coronary artery disease, atrial fibrillation, type 2 diabetes, inflammatory bowel disease, and breast cancer) found that people with top 8.0, 6.1, 3.5, 3.2, and 1.5% highest PRS had a three-fold higher risk to develop these diseases than people with average PRS (Khera et al., 2018).

However, the majority of public genome-wide association study (GWAS) data has been conducted in European population (Popejoy and Fullerton, 2016). Due to the limited availability of non-European ancestral data and the diversity of linkage disequilibrium (LD) architectures among distinct populations, previous studies showed that the genetic architectures of specific phenotypes or diseases were highly consistent between populations (single-variant level and genome-wide level) (Huang et al., 2021). Hence, using PRS derived from European population can result in disease associations being under- or overestimated in other populations (Kim et al., 2018). Traditional approaches are insufficient to address this challenge when multiple populations are involved. Recent genetic statistical studies have indicated that diverse population variants share the same underlying causal variants (Brown et al., 2016; Shi et al., 2020), which raises the possibility of transferability of PRS across distinct ethnic groups. However, existing studies focus mostly on the application with one homogeneous population. For example, LDpred (Vilhjálmsson et al., 2015) and PRS-CS (Ge et al., 2019) improve the prediction accuracy by enhancing LD modelling. As an alternative, a penalized regression framework based on summary statistics, namely Lassosum was proposed by Mak et al. (2017), whereas these methods are limited to GWAS data from one homogeneous population. Current multiethnic PRS construction approaches that incorporate training data from both the European and target populations can leverage trans-ethnic GWAS information and stratify squared trans-ethnic genetic correlation in explanation of environmental effects on genes (Mak et al., 2017; Coram et al., 2017; Shi et al., 2021). Moreover, Márquez-Luna et al. (2017) proposed PT-Multi for multiethnic PRS prediction by performing LD-informed pruning and *P*-value thresholding (PT) (Consortium, 2009) on each homogeneous population and linearly combining the optimal PRS from each specific population.

However, previous studies ignored the information gap among diverse populations. Li et al. (2020) proposed a high-dimensional linear regression model to transfer knowledge between informative samples and target samples to improve the learning performance of target samples. By using GWAS summary statistics from different ancestries and incorporating the idea of transfer learning (Li et al., 2020), we propose a novel statistical method called TL-Multi to enhance the transferability of polygenic risk prediction across diverse populations. TL-Multi assumes most causal variants are shared among diverse populations. There is a difference between the target samples and the informative auxiliary samples in the genetic architecture, which causes estimation bias. TL-Multi further corrects this bias and estimates the PRS using Lassosum (Mak et al., 2017). Additionally, TL-Multi inherits the advantages of Lassosum, ensuring that it has more accurate performance in all circumstances than initial PT and circumvents convex optimization challenges in LDpred. Moreover, TL-Multi extends the application to estimate the genetic risk from unmatched ancestral populations, and employs all available data without pruning or discarding. For practical analysis, we investigate TL-Multi prediction performance with informative auxiliary European samples from UK Biobank (https://www.ukbiobank.ac.uk), and European summary statistics and Hong Kong target samples from previous studies to predict PRS in systemic lupus erythematosus (SLE) (Wang et al., 2021; Morris et al., 2016; Juliá et al., 2018). We obtain a greater than 125% relative improvement in prediction accuracy compared to only using GWAS data from Hong Kong population. Furthermore, TL-Multi performs more accurately in PRS prediction in most scenarios in comparison with the recent multiethnic methods, meta-analysis, and PT-Multi.

Additionally, we refer to Huang et al. (2021) to classify the PRS methods into two categories: single-discovery methods and multi-discovery methods. Single-discovery methods use GWAS data from a single homogeneous population, and multi-discovery methods apply the combined GWAS data of multiple populations.

## 2 MATERIALS AND METHODS

### 2.1 Data Overview

In this study, we requested the individual-level genotyped data for a previous SLE GWAS in Hong Kong (Wang et al., 2021) as the testing dataset, which included 1,604 SLE cases and 3,324 controls. We used GWAS summary statistics of SLE from both East Asian and European populations to train the models. The data for East Asians were collected from Guangzhou (GZ) and Central China (CC), including 2,618 SLE cases and 5,107 controls (Wang et al., 2021). The data for Europeans were obtained from previous studies (Wang et al., 2021; Morris et al., 2016; Juliá et al., 2018), involving a total of 4,576 cases and 8,039 controls. Variants with minor allele frequency greater than 1% and imputed INFO scores greater than 0.7 in respectively ancestral groups were reserved for the following analyses.

In our analysis of asthma, we requested the genotyped data of Indian and European individuals for asthma from UK Biobank. The UK Biobank data consisted of 4,160 unrelated Indian samples genotyped at 1,175,469 SNPs after QC and mapping HapMap 3 SNPs, and we further sampled 48,362 unrelated British samples genotyped at 1,189,752 SNPs after QC and mapping HapMap 3 SNPs. We divided the Indian samples into two groups: 3,160 samples as a training data set and 1,000 samples as a testing data set. As stated previously, the final data set comprises of 3,160 (408 cases and 2,752 controls) unrelated Indian samples for training, 1,000 (127 cases and 873 controls) unrelated Indian samples for testing, and 48,362 (6,555 cases and 41,807 controls) unrelated British samples for training. Variants with minor allele frequency greater than 1% and *P*-values of Hardy-Weinberg equilibrium Fisher’s exact test *<* 1 × 10^−5^ were kept. We then computed GWAS to derive the GWAS summary statistics of each population with each genotypes (after quality control) and adjusting for age and gender, and the top 10 principal components.

### 2.2 Lassosum

Lassosum is a statistical approach introduced by Mak et al. (2017) which enables to tune parameters without validation datasets and phenotype data via pseudovalidation, and outperforms PT and LDpred in prediction (Consortium, 2009; Vilhjálmsson et al., 2015). It refers to the idea of Tibshirani (1996) to deal with sparse matrices and calculate PRS only by using summary statistics and an external LD reference panel. In this article, the ancestry-matched LD block is generally estimated by the 1000 Genome project (https://www.internationalgenome.org). Additionally, we keep the reference panel’s ancestry consistent with that of our target population. Furthermore, if the SNP-wise correlation *r*_*i*_ is not available, we can estimate *r*_*i*_ following Mak et al. (2017): 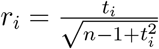.

### 2.3 PT-Multi

PT-Multi assumes the multiethnic PRS is a linear combination of the most predictive PRS from each population. First, it applies LD-pruning and *P*-value thresholding (PT) (Consortium, 2009) to each single ethnic summary statistics and gets the most predictive PRS. Second, it fits marginal linear regression models to get weights for each population, respectively. We apply the R package ‘*bigspnr*’ (Privé et al., 2018) to validation data for LD informed clumping with *r*^2^ threshold of 0.1. The *P*-value thresholds are among: 1, 0.3, 0.1, 3 × 10^−2^, 10^−2^, 3 × 10^−3^, 10^−3^, 3 ×10^−4^, and 10^−4^. We conduct 10-fold cross-validation to determine the optimal *P*-value threshold for each population. We use an independent validation data set to compute the final PRS and the average value of R^2^ across the 10 folds.

This article uses single-discovery method (Lassosum) to regress European, Asian, and multi-discovery methods (meta-analysis, TL-Multi, PT-Multi) to determine the most predictive PRS with the highest R^2^. For ease of notations, let PRS_*a*_, PRS_*e*_, PRS_*ma*_, PRS_*tl*_, and PRS_*pt*_ represent PRS for Asian, European, meta-analysis, TL-Multi and PT-Multi, respectively.

### 2.4 Meta-analysis of two diverse ancestries

We generate the estimates of effect sizes of joint GWAS data by

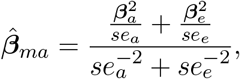

where ***β***_*a*_ and ***β***_*e*_ are the effect sizes obtained from Asian and European GWAS data, respectively, and *se*_*a*_ and *se*_*e*_ are the standard errors obtained directly from ancestry-matched GWAS data. Furthermore, the estimate of the standard error in meta-analysis is defined as:

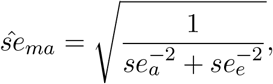

and the estimate of z-statistic is obtained from:

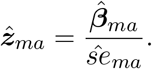

The *P*-value is converted from 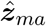 following:

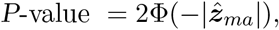

where Φ(·) is the cumulative distribution function of the standard normal distribution *N* (0, 1). In this meta-analysis, the ancestry of the reference panel is consistent with the ancestry of the target population. Furthermore, due to the majority of the total sample being of European ancestry, the LD block is estimated from European population in the 1000 Genome Project.

### 2.5 Multiethnic polygenic risk scores prediction using TL-Multi

In this article, we employ European population data as our informative auxiliary data, owing to its large sample size and relative accessibility. Additionally, we treat East Asians as the target population due to the scarcity of public data (Brown et al., 2016; Shi et al., 2020). Recall the fundamental framework we using for genetic architecture and phenotype, it is a linear combination with effect sizes ***β***, and an *n*-by-*p* genotype matrix ***X***, where *p* is the number of columns containing marker genotype codes corresponding to the number of reference alleles on the sample-specific SNP (e.g., 0, 1, 2), and *n* is the sample size, following as:

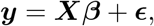

where ***y*** is a vector of clinical outcomes. Tibshirani (1996) proposed Lasso which is commonly used to estimate coefficients 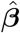 (weights of ***X***), when *p* (the columns of ***X*** or the number of elements of ***y***) is relative large to result in many 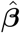 being 0. Specifically, the optimization problem of target population is equivalent to minimizing the objective function:

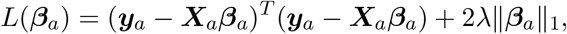

where ***y***_*a*_ is the vector of Aisan phenotypes, ***X***_*a*_ is the genotype matrix of Asian population, *L*(·) is an optimizing function, ‖***β***_*a*_‖_1_ is the *L*_1_ norm of ***β***_*a*_, and *λ* is a data-dependent parameter determining the proportion of ***β***_*a*_ to be estimated to 0.

It can be widely extended in the scenarios in which only the summary statistics are available (Mak et al., 2017).

Motivated by Lassosum, we further propose a novel method, namely TL-Multi to extend its application to multiethnic polygenic prediction. We observed additional samples from auxiliary studies (e.g., European population). The estimate of the marginal effect sizes of European population, 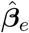, can be generated using the auxiliary model:

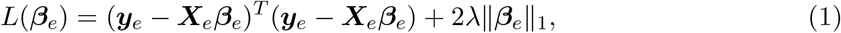

where ***y***_*e*_ is the vector of European phenotypes, and ***X***_*e*_ is the genotype matrix of European population. For illustration, we denote the auxiliary studies, in which informative auxiliary samples can be transferred, and the target model and auxiliary model are similar at certain levels (e.g., similar genetic architectures). Furthermore, we assume the difference between auxiliary samples and target samples is denoted as (Li et al., 2020):

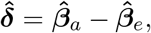

where 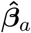 (the weights of target population e.g., Asian population ***X***_*a*_) is the target regression estimator, and 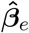 (the weights of auxiliary population e.g., European population ***X***_*e*_) is the estimator for auxiliary study. Furthermore, the informative auxiliary set, *A*_*q*_, has a requirement to ensure that the information auxiliary set includes sufficiently different information under a constrained level. Specifically, the information difference should satisfy the sufficient sparsity:

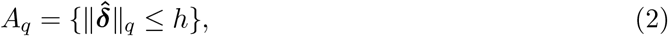

where 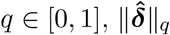 is the *L*_*q*_ norm of the information difference 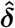 of the informative auxiliary samples. The assumption requires the auxiliary informative population *A*_*q*_ to include samples in their contrast vectors with a maximum *L*_*q*_-sparsity of at most *h*.

Moreover, we assume *A*_*q*_ is informative to improve the prediction performance of target population while *h* is relatively small compared to 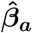. Specifically, when *q* = 0, the set *A*_*q*_ implies that there are at most *h* casual variants. For *q* ∈ (0, 1], this scenario may be explained that all the variants are causal variants with rapid relative amplitudes decaying effect sizes. Therefore, the smaller the *h*, the auxiliary samples of *A*_*q*_ tend to be more informative, where |*A*_*q*_| leverages the number of informative auxiliary samples.

Our goal is to correct the bias between these populations and improve prediction performance in Asian population. First, we can estimate the marginal effect sizes of European population, 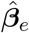 by minimizing the objective function based on equation (1):

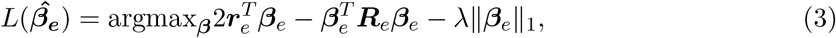

where 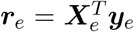 is the SNP-wise correlation between the genotype matrix of European population ***X***_*e*_ and the phenotype ***y***_*e*_, and ***R***_*e*_ is the LD matrix indicating a matrix of correlations between SNPs. The estimates of ***r***_*e*_ can be obtained from summary statistics, and the estimates of ***R***_*e*_ can be obtained from publicly available databases, such as the 1000 Genome project. As Mak et al. (2017) indicated, the PRS can be estimated by optimizing equation (3) without extra individual-level data.

Specifically, TL-Multi estimates the PRS of Asian population by correcting the bias between European and Asian populations. We further denote the bias as ***δ*** which is the difference between European and Asian populations in genetic architecture. The new estimate of effect sizes of Asian population can be presented as: ***β***_*tl*_ = ***β***_*e*_ + ***δ***, in which ***δ*** is estimated by:

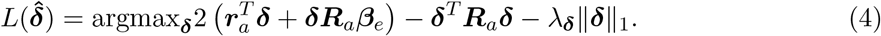

According to pseudovalidation proposed by Mak et al. (2017), the optimal single-discovery PRS for European and Asian populations can be determined directly by the highest R^2^ without the phenotypes. The optimal estimates of effect sizes of Asian and European populations that we apply to TL-Multi are the ancestry-matched optimal PRS, respectively. The Algorithm 1 describes our proposed TL-Multi algorithm, and we further develop an R package, which is publicly available at https://github.com/mxxptian/TL-Multi.git.

#### Algorithm 1

Algorithm for TL-Multi

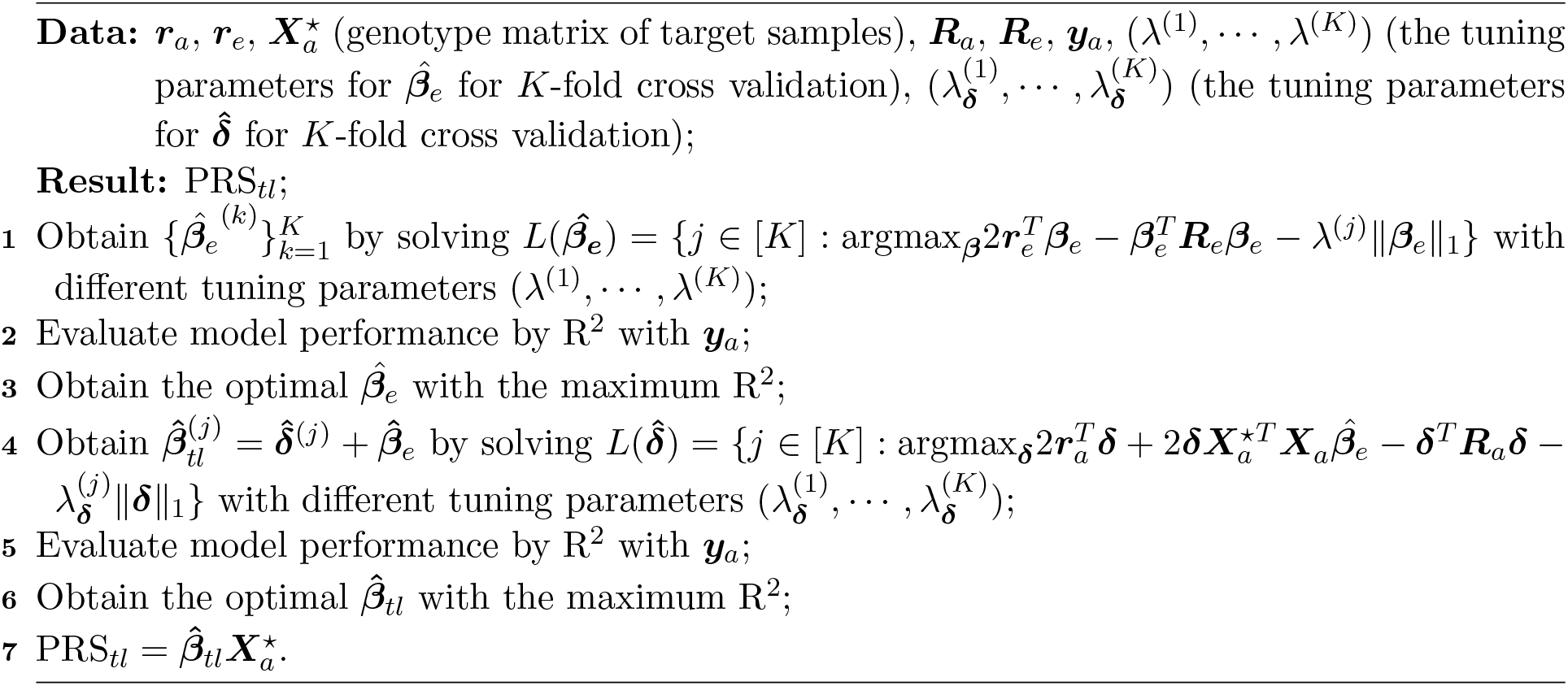

### 2.6 Simulation studies

We performed a wide range of simulation studies to evaluate the performance of TL-Multi. We used real genotypes of European population from UK Biokbank and Asian population from previous SLE study (Wang et al., 2021).

Following the quality control procedure provided in Chang et al. (2015), we utilized the UK Biobank and Asian lupus genotype data whose *P*-values of Hardy–Weinberg equilibrium Fisher’s exact test *<* 1 × 10^−5^ with minor allele frequency (MAF) *>* 1% and filtered out SNPs and missing samples. Then, we simulated the effect sizes based on the genetic architecture correlation and applied the R package ‘bigsnpr’(Privé et al., 2018) to generate quantitative phenotypes and conduct GWAS to determine the summary statistics. Based on these estimated summary statistics, we employed the following PRS prediction methods. We further extracted the common variants between European samples and Asian samples. This resulted in 69, 398 SNPs in total, and 4, 049 subjects in Asian population. We fixed SNP-heritability *h*^2^ at 0.5, and further simulated genetic architectures by randomly treating 1%, 1.5%, 2%, and 5% variants as causal variants. We assumed that these causal variants were shared in multiple populations with different effect sizes. Additionally, we sampled effect sizes from a multivariate normal distributio with a wide range of cross-population genetic correlation values (0.2, 0.4, 0.6, and 0.8) (Huang et al., 2021; Bulik-Sullivan et al., 2015), where for each population the variance is 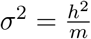 and *m* is the number of causal variants. There were 12 combinations in total. For each scenario, we generated 20 replicates and calculated the average values to assess the prediction accuracy. We took out the original phenotypes and generated new ones based on a linear framework:

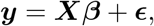

where ***X*** is the training set of the standardized genotype matrix, and ***ϵ*** represent the random error which was generated from *N* (0, 1 − *h*^2^).And GWAS was implemented using the R package ‘bigsnpr’ to obtain the summary statistics for each simulated phenotype.

Due to the possibility that sample size affects performance, we investigated 25:1 and 50:1 proportions of European samples to Asian samples. Additionally, we observed that the number of variants has a significant influence on the prediction performance, and the majority of variants are located on chromosomes 1-11. Motivated by previous works (Vilhjálmsson et al., 2015; Márquez-Luna et al., 2017), we further extrapolated the performance at large sample size by conducting simulations with different subsets of chromosomes to increase 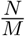, where *N* is the total number of samples and *M* is the number of SNPs: (1) using chromosomes 1-4; (2) using chromosomes 1-6; (3) using chromosomes 1-8; (4) using chromosomes 1-11.

## 3 RESULTS

### 3.1 Simulations

We performed simulations with real genotypes and simulated continuous phenotypes. We split the data from Hong Kong population into two groups: 1,000 samples as a training data set and 3,049 samples as testing data, and drew 50,000 samples from European samples. The training data set was used to simulate phenotypes, and the testing data were applied to performance assessments. The prediction accuracy was assessed by R^2^, which was based on the simulated phenotypes generated from the test data. Specifically, LD blocks for single-discovery method were ancestry-matched as the reference panels, and they were in correspondence with the ancestry of the target population for multi-discovery methods.

In Figure 1, we displayed the average values with a 95% upper bound of each simulation setting under scenario (1) over 20 replicates. We conducted single-discovery analyses for Asian and European populations by Lassosum, and multi-discovery analyses by meta-analysis, TL-Multi, and PT-Multi. Lassosum adopted the PRS with the maximum R^2^ by 10-fold cross-validation. We observed that meta-analysis could not improve the prediction accuracy when single-discovery analysis of European population did not perform better than the Asian one. Particularly, when the genetic architecture correlation was quite low (*ρ* = 0.2), meta-analysis and European prediction performances were comparably inferior. In this case, it was explained that the shared information between the Asian and European populations would be limited, preventing prediction improvement from being achieved by directly integrating the European data. It also reflected the consistent relationship between meta-analysis and single-discovery analysis of the informative population. Moreover, meta-analysis could hardly outperform the European one. The performance of Lassosum for European population dominated the performance of meta-analysis since the sample size of European population is significantly larger than that of Hong Kong. Additionally, we observed that TL-Multi could always improve the accuracy compared to Lassosum for Hong Kong population. If the genetic architecture correlation was not too high (e.g., *ρ* = 0.4 or 0.6), TL-Multi attained the highest prediction accuracy compared to competing approaches. However, when the genetic architecture was high (*ρ* = 0.8), we noticed that TL-Multi performed slightly worse than other approaches. In this example, the results might be explained by the remarkable similarity of the genetic architecture. When the genetic correlation reaches 0.8, the majority of information about the Asian population would be directly explained by that of the European population. Combining these two groups in the meta-analysis might increase the accuracy of estimated effect sizes.

**Figure 1:**
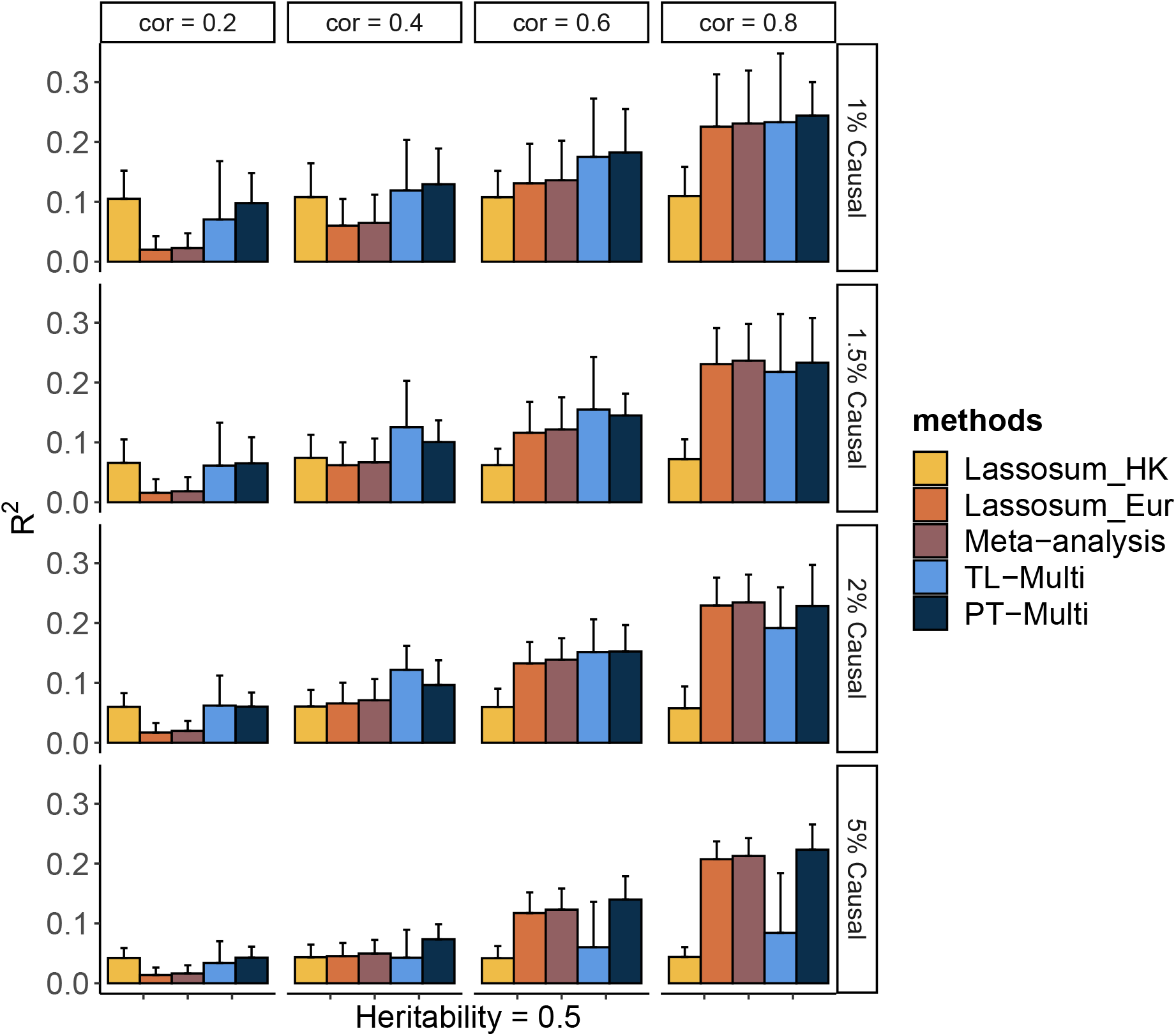
Prediction accuracy of Lassosum, meta-analysis, PT-Multi, and TL-Multi over 20 replications in simulations. Lassosum_HK is Lassosum for Hong Kong population, and Lassosum Eur is Lassosum for European populations. Heritability was fixed at 0.5 and different genetic correlations (0.2, 0.4, 0.6, and 0.8) with different causal variant proportions (1%, 1.5%, 2%, and 5%) were generated. 50,000 European samples and 1,000 Hong Kong samples were simulated. The variants were generated from the common variants of the first 4 chromosomes (21,477 SNPs). The prediction accuracy was measured by R^2^ between the simulated and true phenotypes. The error bar indicated the upper bound of 95% confidence interval over 20 replications.

In most scenarios, TL-Multi outperformed PT-Multi. Specifically, TL-Multi substantially improved multiethnic prediction accuracy for the instances with 1%, 1.5%, 2% causal proportions. PT-Multi conducted PT, which caused information loss in the data. However, TL-Multi could take all the data information into account. We found TL-Multi performed poorly at a 5% causal proportion. We noted that under this situation, the result of Lassosum for Hong Kong population was significantly inferior to that of European. We referred to the assumption (2) to cast doubt on the breach of our assumption. If the assumption does not hold, European population could not be denoted as auxiliary informative data because the useful information was limited. Due to it, TL-Multi would fail to borrow the information to improve the learning performance of the target population. Alternatively, consider that the effect sizes were simulated depending on the number of causal variations, *m*. As the proportion of causality rose, the effect sizes tended to approach zero. Limited by small sample size of the Asian population, the bias between the estimated effect sizes derived from the simulated phenotypes and the actual effect sizes would be even larger. Some causal variants with relative small signals more likely erroneously failed to be captured which resulted in the restricted TL-Multi’s performance. However, it is noteworthy that TL-Multi still enhanced Hong Kong’s prediction accuracy in this scenario. We discovered that the performance of meta-analysis and PT-Multi for Hong Kong were nearly identical to that of Lassosum for Europeans, when we attributed to the huge disparity in multiethnic sample sizes. To summarize, European population dominated the performance of meta-analysis and PT-Multi. In particular, TL-Multi could be employed to the moderate genetic architecture correlations (e.g., *ρ* = 0.4, and 0.6) when the informative auxiliary population (e.g., European population) outperformed the target population (e.g., Hong Kong population). Referring to the assumption (2), the performance of European population was supposed to be more accurate than that of the target population, therefore it would be appropriate to borrow information from it. Moreover, if the proportion of casuals increased, the estimated effect sizes of the target population would be relatively biased. We found that the precision of the effect sizes of the target population would have a substantial effect on TL-Multi.

Alternatively, we generated phenotypes using different chromosome subsets and sample sizes of European population while maintaining a fixed Hong Kong sample size. Over 20 replicates, we took the performance using a fixed genetic correlation of 0.4 and 1.5% causal variants as an example. In Figure 2(A), we drew 25,000 European subjects and 1,000 Hong Kong subjects. We observed that TL-Multi performed much better than the competing approaches. While the performance of Hong Kong was superior to that of Europeans, the performance of the meta-analysis was poor compared to that of Hong Kong. As the total number of SNPs increased, the prediction accuracy of Hong Kong dramatically decreased. However, the prediction accuracy of Europeans decreased relatively slowly. Specifically, under scenario (4), TL-Multi was inferior to the other two multi-discovery methods. This could be explained that for this case, there were 1,000 subjects from Hong Kong with 49,909 SNPs which resulted in a significant bias while estimating the effect sizes by applying GWAS. In this case, TL-Multi thus failed to improve the accuracy of the forecast compared to the previous scenarios, as the bias in the estimates of Hong Kong’s effect sizes was larger.

**Figure 2:**
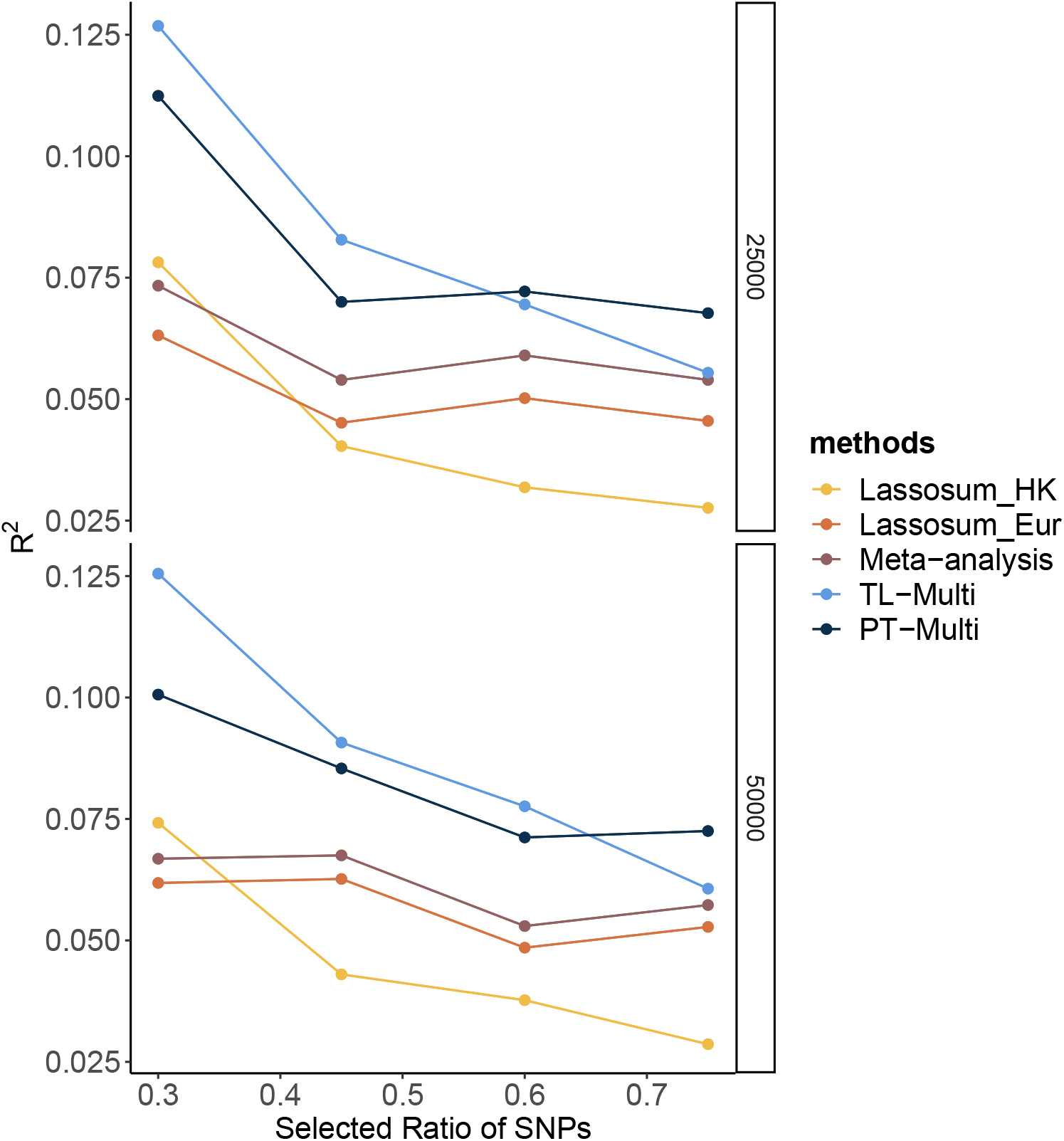
Prediction accuracy of Lassosum, meta-analysis, PT-Multi and TL-Multi over 20 replications in simulations. Selected ratio of SNPs is the ratio of the actual numbers of SNPs simulated to the total number of common SNPs (69,398). The actual numbers of SNPs simulated in the four scenarios are 21,477 (chromosomes 1-4), 32,151 (chromosomes 1-6), 39,682 (chromosomes 1-8), 49,909 (chromosomes 1-11) respectively. The average of R^2^ are plotted. (A) The sample size of European population is 25,000, and the sample size of Hong Kong population is 1,000. (B) The sample size of European population is 50,000, and the sample size of Hong Kong population is 1,000.

Moreover, the consistent trend in European, meta-analysis, and PT-Multi supported our previous extrapolation that the performance of European population could determine the primary contribution of the other two. In Figure 2(B), we simulated 50,000 European subjects. We further observed that the performance of PT-Multi was inferior to TL-Multi under the scenarios (1)-(3) and both of them outperformed the single-discovery method and meta-analysis. Furthermore, the performance of meta-analysis was consistent with that of European. As a result, even though the prediction accuracy of TL-Multi went down, it was still better than the meta-analysis’s prediction accuracy under all the scenarios.

### 3.2 Analysis of SLE in Hong Kong Population

We further applied the above four approaches to predict SLE risk in Hong Kong population to evaluate the performance in real data analysis. We used European SLE GWAS summary statistics from previous studies (Wang et al., 2021; Morris et al., 2016; Juliá et al., 2018) (4,576 cases, and 8,039 controls), and the ancestry-matched GWAS summary statistics (Wang et al., 2021) (2,618 cases, and 5,107 controls). The validation data for Hong Kong population were from Wang et al. (2021) (1,604 cases, and 3,324 controls) employing 10-fold cross-validation following Mak et al. (2017).

We reported the area under the receiver operating characteristic curve (AUC) to assess the prediction accuracy of derived PRS. The ethnicity of the LD block is consistent with that of the majority population in GWAS data, and the LD block was derived from Berisa and Pickrell (2016). Furthermore, the reference panel was obtained from the 1000 Genome Project, and the ethnicity of it was consistent with the target population’s. We set the *P*-value thresholds to be the same as the values in simulation studies, and *r*^2^ = 0.1. In real data analysis, TL-Multi outperformed the competing methods. The optimal PRS from European GWAS data yielded AUC of 0.6872 and 0.6943 from East Asian GWAS data. We further obtained the optimal PRS of meta-analysis, TL-Multi and PT-Multi, with AUC values of 0.7098, 0.7131, and 0.5447, and the corresponding ROC curves were depicted in Figure 3. For binary classification, we used a logistic regression to obtain the mixing weights in PT-Multi. Consistent with the evaluations in simulation studies, we observed that TL-Multi improved 2.7% in prediction accuracy compared to Lassosum for Hong Kong population, and meta-analysis improved 2.2% compared to Lassosum. However, PT-Multi performed even worse than single-discovery method in real data analysis.

**Figure 3:**
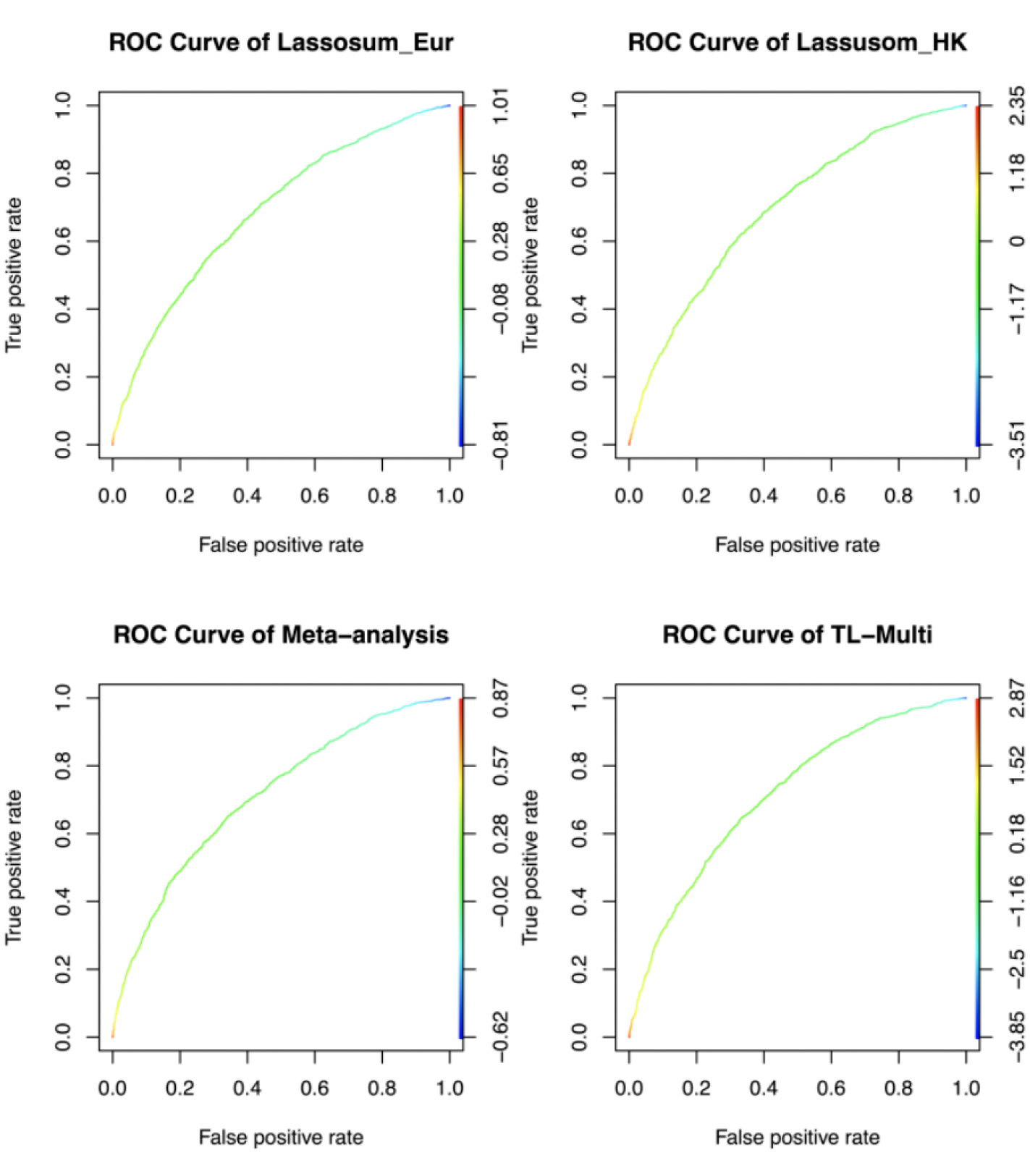
Receiver operating characteristic curve of Lassosum, meta-analysis, and TL-Multi in analysis of SLE study. Lassosum_HK is Lassosum for Hong Kong population, and Lassosum Eur is Lassosum for European population. The corresponding AUC values with the optimal PRS of Lassosum for Hong Kong population and European population, meta-analysis, and TL-Multi are 0.6872, 0.6943, 0.7098 and 0.7131, respectively.

Moreover, we reported the case prevalence of the bottom 2%, 5%, and 10% and top 2%, 5%, and 10% of PRS distribution, constructed by single-discovery method, meta-analysis, and TL-Multi in Table 1. This summary report demonstrated the case prevalence under different PRS conditions. For instance, the bottom numbers indicate the prevalence of SLE among individuals with low PRS. We observed that TL-Multi had satisfactory performance and showed 10.66, 7.50, and 5.80 fold increases comparing the top 2%, 5%, and 10% with bottom 2%, 5%, and 10% of the PRS distribution, respectively.

**Table 1:** Case prevalence of 2%, 5%, and 10% for the top and bottom quantiles of the PRS distribution in analysis of SLE study with the target Indian population, generated by Lassosum, meta-analysis, and TL-Multi.

| Prevalence | Bottom |  |  | Top |  |  |
| --- | --- | --- | --- | --- | --- | --- |
|  | 2% | 5% | 10% | 10% | 5% | 2% |
| Lassosum_HK | 0.0864 | 0.1133 | 0.1309 | 0.6963 | 0.7192 | 0.7407 |
| Lassosum_Eur | 0.1111 | 0.1281 | 0.1704 | 0.6938 | 0.7389 | 0.8148 |
| Meta-Analysis | 0.0864 | 0.0985 | 0.1309 | 0.7432 | 0.8030 | 0.8519 |
| TL-Multi | 0.0741 | 0.0985 | 0.1235 | 0.7160 | 0.7389 | 0.7901 |

### 3.3 Analysis of Asthma in Indian Population

We applied the same methods to the Indian and European samples from UK Biobank computing associated summary statistics by ‘bigsnpr’ R package. We splitted 1,000 (127 cases and 873 controls) unrelated Indian samples as validation data, and 3,160 (408 cases and 2,752 controls) unrelated samples as training data, and further sampled 48,362 (6,555 cases and 41,807 controls) unrelated European samples. We further reported the AUC of the above four methods to evaluate the optimal prediction method. The ancestry of LD blocks matches to that of the data’s predominant population. We used training data as a reference panel whose ancestry was always identical to that of the target population. During pruning and clumping, the *P*-value thresholds were set to be equal to simulation with *r*^2^ = 0.1.

The ROC curves for binary classification are depicted in Figure 4. The optimal PRS from European and Indian samples revealed AUC values of 0.5657 and 0.5441, respectively. In addition, for the multiethnic PRS construction methods, the optimal PRS of meta-analysis, TL-Multi, and PT-Multi resulted in AUC values of 0.5705, 0.5721, and 0.6427, respectively. We found that TL-Multi was superior to the all singe-discovery methods and meta-analysis. For binary classification, TL-Multi improved 5.15% in prediction accuracy compared to Lassosum for Indian population, and meta-analysis improved 4.85% compared Lassosum for Indian population. We noted that PT-Multi performed better than ours. However, the comparison of PT-Multi method with other methods might not be fair since PT-Multi required individual level data whereas other four approaches solely relied on the summary statistics. Moreover, access to individual-level data was typically difficult.

**Figure 4:**
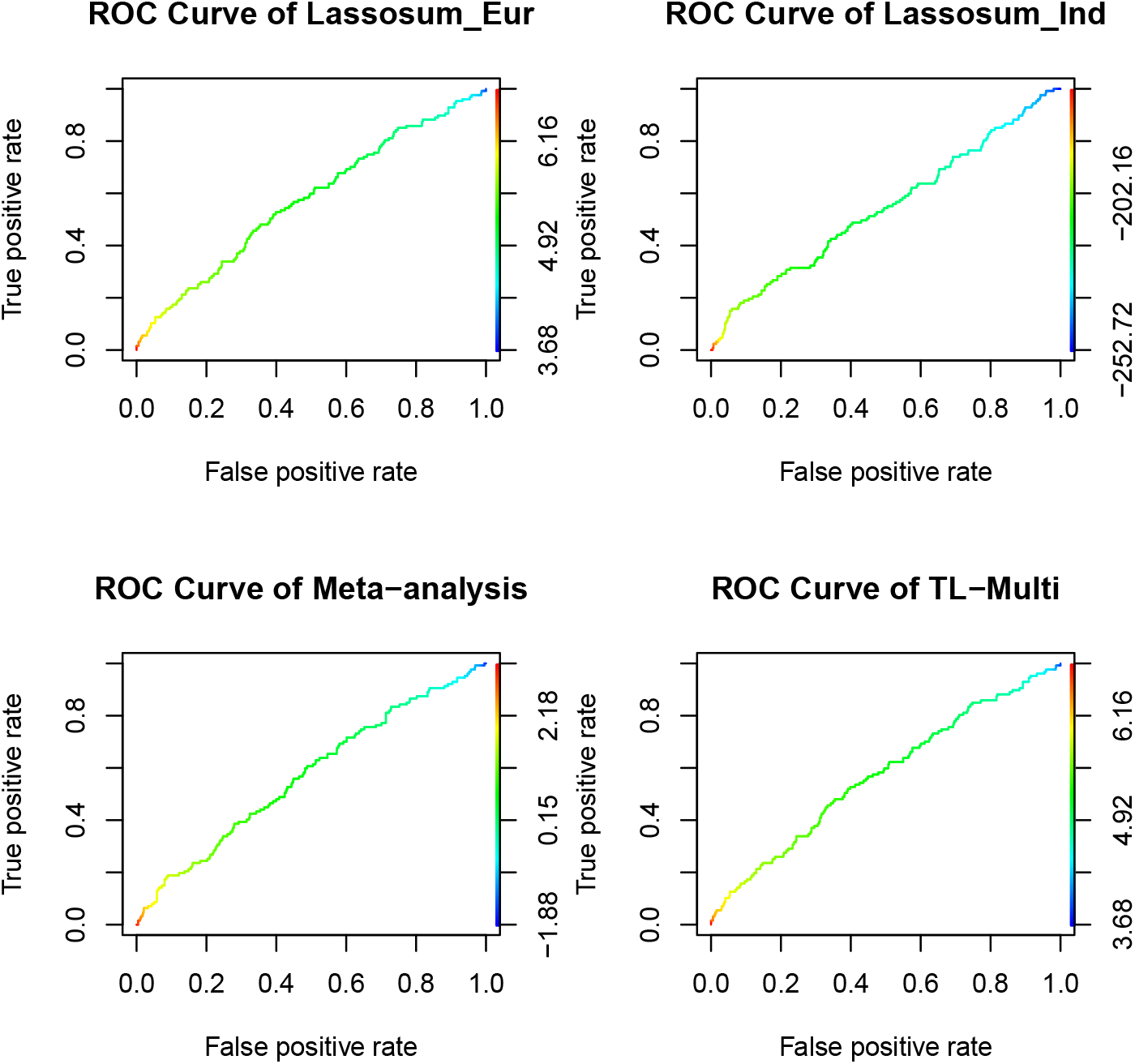
Receiver operating characteristic curve of Lassosum, meta-analysis, and TL-Multi in the analysis asthma study. Lassosum_Ind is Lassosum for Indian population, and Lassosum Eur is Lassosum for European population. The corresponding AUC values with the optimal PRS of Lassosum for European population and Indian population, meta-analysis, and TL-Multi are 0.5657, 0.5441, 0.5705 and 0.5721, respectively.

Additionally, the case prevalence of the bottom 2%, 5%, and 10% and top 2%, 5%, and 10% of PRS distribution, conducted by Lassosum for Indian and European, mata-analysis and TL-Multi was reported in Table 2. We observed that TL-Multi would also perform with more accuracy in terms of case prevalence than the competing methods.

**Table 2:** Case prevalence of 2%, 5%, and 10% for the top and bottom quantiles of the PRS distribution in analysis of asthma study with the target Indian population, generated by Lassosum, meta-analysis, and TL-Multi.

| Prevalence | Bottom |  |  | Upper |  |  |
| --- | --- | --- | --- | --- | --- | --- |
|  | 2% | 5% | 10% | 10% | 5% | 2% |
| Lassosum_Ind | 0.1500 | 0.1800 | 0.1600 | 0.1900 | 0.1800 | 0.2000 |
| Lassosum_Eur | 0.1500 | 0.1200 | 0.1300 | 0.1600 | 0.1800 | 0.0000 |
| Meta-Analysis | 0.4000 | 0.2000 | 0.2100 | 0.1400 | 0.1600 | 0.100 |
| TL-Multi | 0.1000 | 0.1800 | 0.1700 | 0.2000 | 0.2400 | 0.2500 |

## 4 DISCUSSION

In this article, we have proposed a novel approach named TL-Multi to improve the accuracy of PRS prediction for non-European populations. Our proposed method leverages summary statistics and makes complete use of all available data without clumping. We have shown that transferring the information from the informative auxiliary populations (e.g., European) to the target populations (e.g., East Asian) can indeed improve learning performance and the prediction accuracy of the target populations compared to the single-discovery methods. Particularly, TL-Multi shows a higher AUC compared to meta-analysis and PT-Multi in analysis of SLE in Hong Kong population. In our analysis of asthma in the Indian population, TL-Multi out-performs Lassosum and meta-analysis in terms of prediction performance and case prevalence prediction accuracy. Moreover we note that in the field of PRS prediction, there is no a particular method outperforms all the others. It depends on the specific situation to select an appropriate method. For instance, PRS-CS can always outperform PT in Huang et al. (2021), but PRS-CS may be inferior to PT in Weissbrod et al. (2022) in some circumstances. Therefore, we provide some potential circumstances in which TL-Multi would be an appropriate choice. First, TL-Multi is implemented using summary statistics and performs well under the moderate genetic architecture correlation (e.g., *ρ* = 0.4 and 0.6) and moderate causal proportions (e.g., 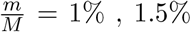, and 2%). Second, based on the assumption (2), TL-Multi would be a good alternative when the single-discovery method’s performance of the informative auxiliary population is superior to that of the target population.

Compared to the single-discovery methods, we showed that the performance of TL-Multi was always more accurate with an acceptable running time (e.g., 2 minutes) than the performance of Lassosum for Hong Kong population, especially under moderate genetic correlation (e.g., *ρ* = 0.6). When the sample size of the target data set is limited, increasing the sample size of the informative data set can enhance the prediction accuracy of TL-Multi. In the simulation studies, we found that the performances of meta-analysis and PT-Multi were dominated by the performance of Lassosum for European population. As the genetic architecture correlation was rather high (*ρ* = 0.8), TL-Multi may perform poorly, and it would be more prudent to consider approaches that integrate the whole data set across distinct populations. Therefore, the performances of PT-Multi and meta-analysis were unsatisfactory, while the performance of European population was worse than that of Hong Kong population.

Another advantage of TL-Multi is its powerful transferability, which corrects the bias in estimation between European and non-European populations. De Candia et al. (2013) showed that the cross-population genetic correlation could leverage the causal effect sizes in different populations. In simulation studies, TL-Multi performed better when the genetic correlations were 0.4 and 0.6. It indicated that TL-Multi could be widely applied to two different populations which share some common genetic architecture information. Moreover, TL-Multi retained the pseudovalidation proposed in Mak et al. (2017). It extended the application of TL-Multi to fit the data without a validation data set and phenotype data.

Despite these advantages, some limitations of TL-Multi still remain for the future work. For example, if the difference between two populations is too enormous, our proposed approach’s assumptions will fail to hold. It is worth bearing in mind to deal with this scenario. And in this article, we did not consider the X chromosome, whose information could also contribute to prediction accuracy (Tukiainen et al., 2014). In recent years, some approaches have fitted multiple diseases simultaneously (Maier et al., 2015; Turley et al., 2018; Chung et al., 2019; Musliner et al., 2019; Graff et al., 2021). These studies inspire us to investigate other TL-Mutli extensionsthat bridge not only the gap between populations but also the gap between illnesses in the interim.

## Data Availability Statement

The 1000 Genome project data can be found in the https://www.internationalgenome.org/. The UK Biobank data were accessed under Application Number 58942. Inquiries about the study’s SLE summary association statistics can be directed to the corresponding author.

## Notes

### Competing Interest Statement

The authors have declared no competing interest.

### Summary of Updates

Correct some typos and the order of the Tables

